# A large-scale serological survey in pets from October 2020 through June 2021 in France shows significantly higher exposure to SARS-CoV-2 in cats

**DOI:** 10.1101/2022.12.23.521567

**Authors:** Matthieu Fritz, Eric Elguero, Pierre Becquart, Daphné de Riols de Fonclare, Déborah Garcia, Stephanie Beurlet, Solène Denolly, Bertrand Boson, Serge G. Rosolen, François-Loïc Cosset, Alexandra Briend-Marchal, Vincent Legros, Eric M. Leroy

## Abstract

Severe acute respiratory syndrome coronavirus 2 (SARS-CoV-2) can infect many animals, including pets such as dogs and cats. Many studies have documented infection in companion animals by bio-molecular and serological methods. However, only a few have compared seroprevalence in cats and dogs from the general population, and these studies were limited by small sample sizes and collections over short periods. Our goal was to obtain a more accurate evaluation of seroprevalence in companion animals in France and to determine whether cats and dogs differ in their exposure to SARS-CoV-2. For this purpose, we conducted an extensive SARS-CoV-2 cross-sectional serological survey of 2036 cats and 3577 dogs sampled by veterinarians during medical examinations in clinics throughout France. Sampling was carried out from October 2020 through June 2021, a period encompassing the second and third waves of SARS-CoV-2 infections in humans in the country. Using a microsphere immunoassay targeting the receptor binding domain and trimeric spike protein, we found 7.1% seroprevalence in pets. In a subset of 308 seropositive samples, 26.3% had neutralizing antibodies. We found that cats were significantly more likely to test positive than dogs, with seropositivity rates of 9.3% and 5.9% in cats and dogs, respectively. Finally, data for both species showed that seroprevalence was lower in older animals and was not associated with the date of sampling or the sex of the animal. Our results show that cats are significantly more sensitive to SARS-CoV-2 than dogs, in line with experimental studies. Our large sample size provides for a reliable, statistically robust estimate of the frequency of infection of pets from their owners and offers strong support for the notion that cats are more sensitive to SARS-CoV-2 than dogs. Our findings emphasise the importance of a One-Health approach to the SARS-CoV-2 pandemic and raise the question of whether companion animals in close contact with humans should be vaccinated.

## Introduction

Two months after the onset of SARS-CoV-2 circulation in humans, two dogs in Hong Kong were reported to have naturally acquired the virus (1). Since then, many studies have reported viral RNA and SARS-CoV-2 antibodies in dogs and cats—mostly belonging to COVID-19-infected owners (2-5). Furthermore, several studies demonstrated that the risk of pets testing seropositive was higher in COVID-19+ households than for pets from households of unknown status (6-10).

Definitive examples of pet-to-human transmission are scarce. A recent study from Thailand reported a suspected case of SARS-CoV-2 transmission from a cat to a human (11), and dog-to-human transmission has yet to be described. However, given that 200 million cats and dogs live in close proximity to humans in Europe (12), there is ample opportunity for such transmission, and the potential risks need to be carefully considered.

Several population-based serological studies have reported SARS-CoV-2 antibodies in dogs and cats. In dogs, estimates of seroprevalence have ranged from 0% to 14.5% (6, 10, 13-21). While in cats, estimates have ranged from 0% to 21.7% (6, 14, 15, 20, 22-25). For both species, seroprevalence was highly dependent on the period of sampling (first, second wave *etc*.), the assay used (ELISA, seroneutralization, etc.), and the country of sampling (China, Croatia, Germany, Italy, the Netherlands, Poland, Portugal, Spain, Thailand, United-Kingdom, USA). Among these studies, five directly compared cats and dogs. There is some experimental and epidemiological evidence suggesting that cats are more susceptible to infection than dogs (6, 7, 15, 26, 27). However, significant species differences have not always been observed in population-based studies (13, 17, 19). This is perhaps because of significant study limitations—a low number of enrolled animals, a short sampling period, etc.—that have curtailed robust estimates of infection rate in pets with enough statistical power to recognize differences in COVID-19 epidemiology.

Here we report estimates of the frequency of SARS-CoV-2 infection from 2036 cats and 3577 dogs sampled at veterinary clinics from October 2020 through June 2021 throughout France—the largest serological survey of SARS-CoV-2 infections in companion animals to date. The study allows for robust estimates of pet infection rate and provides strong support for the hypothesis that species differences in susceptibility observed in experimental studies translate into a significant increase in infection rate in cats.

## Materials and Methods

### Sampling

The cross-sectional, nationwide sampling was possible thanks to a network of veterinary clinics across France working with VEBIO. VEBIO is a veterinary diagnostic laboratory which performed all categories of medical analyses, including infectious diseases, hematology, endocrinology, oncology … (see more details in https://www.vebio.fr/). No inclusion and exclusion criteria were applied for the selection of blood samples, except that they came only from veterinary clinics working with VEBIO. VEBIO notified the veterinary clinics that following requested biomedical analyses, the remaining serum could be used in a SARS-CoV-2 research project. No specific request for samples was addressed to the vets. Thus, the SARS-CoV-2 analysis is based on samples collected during the regular activities of the vets.

Blood samples were collected in dry/EDTA tubes from dogs and cats during routine healthcare visits or for diagnostic purposes at veterinary clinics. After centrifugation, the serum/plasma was kept at +4 °C until sent to VEBIO. Rapid and safe shipping practices were used to avoid contamination and ensure samples reached VEBIO within 48 h. At the VEBIO facility, an aliquot was taken from the sample to perform the requested biomedical analyses. Another aliquot was then stored at +4 °C until sent to the MIVEGEC lab, Montpellier, where serological analyses were performed. Safe shipping practices with an approved professional carrier were also used for shipment to the MIVEGEC lab. Finally, the samples were stored at the MIVEGEC lab at −20 °C until testing (Figure 1). For shipping to the CIRI lab, SARS-CoV-2-positive samples detected by MIA were transported by an approved professional carrier at -20°C to ensure optimal safety conditions. Data (age, sex, clinical history, and region localization, when available) from dogs and cats were provided anonymized by VEBIO to the MIVEGEC lab.

**Figure 1.**
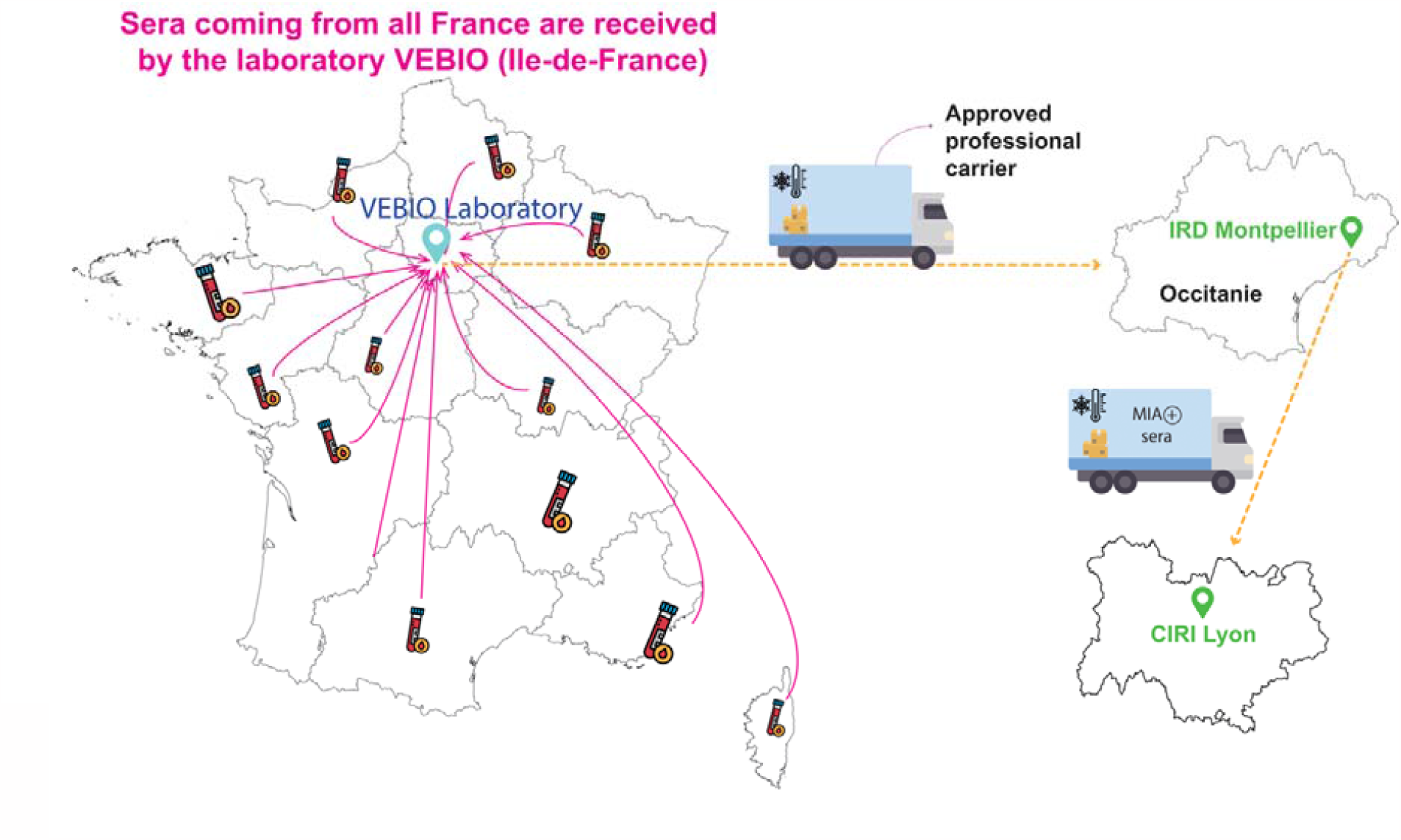
Logistics of sample collection and distribution. Sera collected during routine healthcare visits by veterinarians throughout France were first sent to VEBIO in Ile-de-France. Aliquots of the samples were then made and sent to the IRD in Montpellier (Hérault) via an approved carrier.

### Ethics

According to the act governing the “use of live animals for scientific purposes” effective in France on 14 January 2022, ethical approval was not sought or required since all pets were sampled by a veterinarian during a health care visit. All applicable international and national guidelines for the care of pets were followed.

### Microsphere Immunoassay (MIA)

Dog and cat serum samples were tested using a multiplex microsphere immunoassay (MIA). Ten μg of two recombinant SARS-CoV-2 antigens, receptor-binding domain (RBD) and trimeric spike (tri-S), both derived from the Whuhan-Hu-1 strain (The Native Antigen Company, Kidlington United-Kingdom), were used to capture specific serum antibodies. Distinct MagPlex microsphere sets (Luminex Corp, Austin, TX, USA) were respectively coupled to viral antigens using the amine coupling kit (Bio-Rad Laboratories, Marnes-la-Coquette, France) according to the manufacturer’s instructions. Microsphere mixtures were successively incubated with serum samples (1:400), biotinylated protein A and biotinylated protein G (4 μg/mL each) (Thermo Fisher Scientific, Illkirch, France), and streptavidin-R-phycoerythrin (4 μg/mL) (Life technologies, Illkirch, France) on an orbital shaker and protected from light. Measurements were performed using a Luminex 200 instrument (Luminex Corp, Austin, TX, USA), and at least 100 events were read for each bead set. Binding events were displayed as median fluorescence intensities (MFI). Specific seropositivity cut-off values for each antigen were set at three standard deviations above the mean MFI of pre-pandemic serum from 53 dogs and 30 cats sampled before 2019. These samples were stored in biobanks at the IRD and VetAgro Sup. MIA specificity was set for each antigen at 96.2% for dogs and 100% for cats based on the pre-pandemic populations. MIA was first validated using sera from two COVID-19 PCR+ humans, kindly provided by Meriadeg Ar Gouilh, and then with sera from SARS-CoV-2 PCR+ cats and dogs, provided by several veterinarians.

Because of the excellent specificity observed for both antigens and to account for any isotypic variability, an animal was deemed positive for SARS-CoV-2 antibodies following a positive result in at least one of the two tests.

### Neutralization activity measurement

An MLV-based pseudoparticle carrying a GFP reporter pseudotyped with SARS-CoV-2 spike protein (Wuhan-Hu-1 strain) (SARS-CoV-2pp) was used to measure neutralizing antibody activity in cat and dog sera. Each SARS-CoV-2-positive sample detected by MIA was processed according to a neutralization procedure as previously described (28) . Briefly, for neutralization assays, a sample of ∼1⍰1⍰× ⍰10^3^ pseudoparticles was incubated with a 100-fold dilution of sera or control antibodies for 1⍰h at 37⍰°C before infection of Vero-E6R cells. At 72⍰h post-transduction, the percentage of GFP-positive cells was determined by flow cytometry (at least 10 000 events recorded). The level of infectivity is expressed as the percentage of GFP-positive cells and compared to cells infected with SARS-CoV-2pp incubated without serum. As a control, the same procedure was done using RD114 pseudoparticles to identify sera with aspecific neutralization. Sera exhibiting more than 30% SARS-CoV-2pp neutralization were considered positive. Pre-pandemic serum from France was used as a negative control, and an anti-SARS-CoV-2 RBD antibody was used as a positive control.

### Statistical analyses

Associations between SARS-CoV2 infection status (positive or negative) and the covariates region, age, and sex were assessed using binomial (logistic) generalized linear models. The region was defined by where the animal lived at the time of sampling. Age was that recorded by the veterinarian, with variable precision, generally in months for young animals and whole years for older animals. Its accuracy is unknown. The associated statistical tests were likelihood ratio tests. All analyses were performed using R software (29).

## Results

### Blood collection

Blood samples from 2036 cats and 3577 dogs were collected during routine healthcare visits by veterinarians from October 2020 through June 2021 (Table 1). Samples came from all 13 regions of metropolitan France. Corsica was excluded due to too few samples. Almost half of the samples came from Ile-de-France, the region including Paris, reflecting population density and proximity to the Veterinary diagnostic laboratory (VEBIO), where all samples for biomedical analyses requested by the veterinarians were handled (Materials and Methods). The number of samples received from other regions largely depended on the number of veterinarians working with VEBIO in those regions (Figure 2). Unfortunately, we could not study the clinical history of the animals due to variability in how each veterinarian reported this information.

**Table 1:** numbers of samples collected by month and by species.

|  | October | November | December | January | February | March | April | May | June |
| --- | --- | --- | --- | --- | --- | --- | --- | --- | --- |
|  | 2020 | 2020 | 2023 | 2021 | 2021 | 2021 | 2021 | 2021 | 2021 |
| Cats | 49 | 256 | 275 | 291 | 225 | 296 | 305 | 166 | 173 |
| Dogs | 84 | 428 | 543 | 475 | 403 | 474 | 597 | 282 | 291 |
| Total | 133 | 684 | 818 | 766 | 628 | 770 | 902 | 448 | 464 |

**Figure 2.**
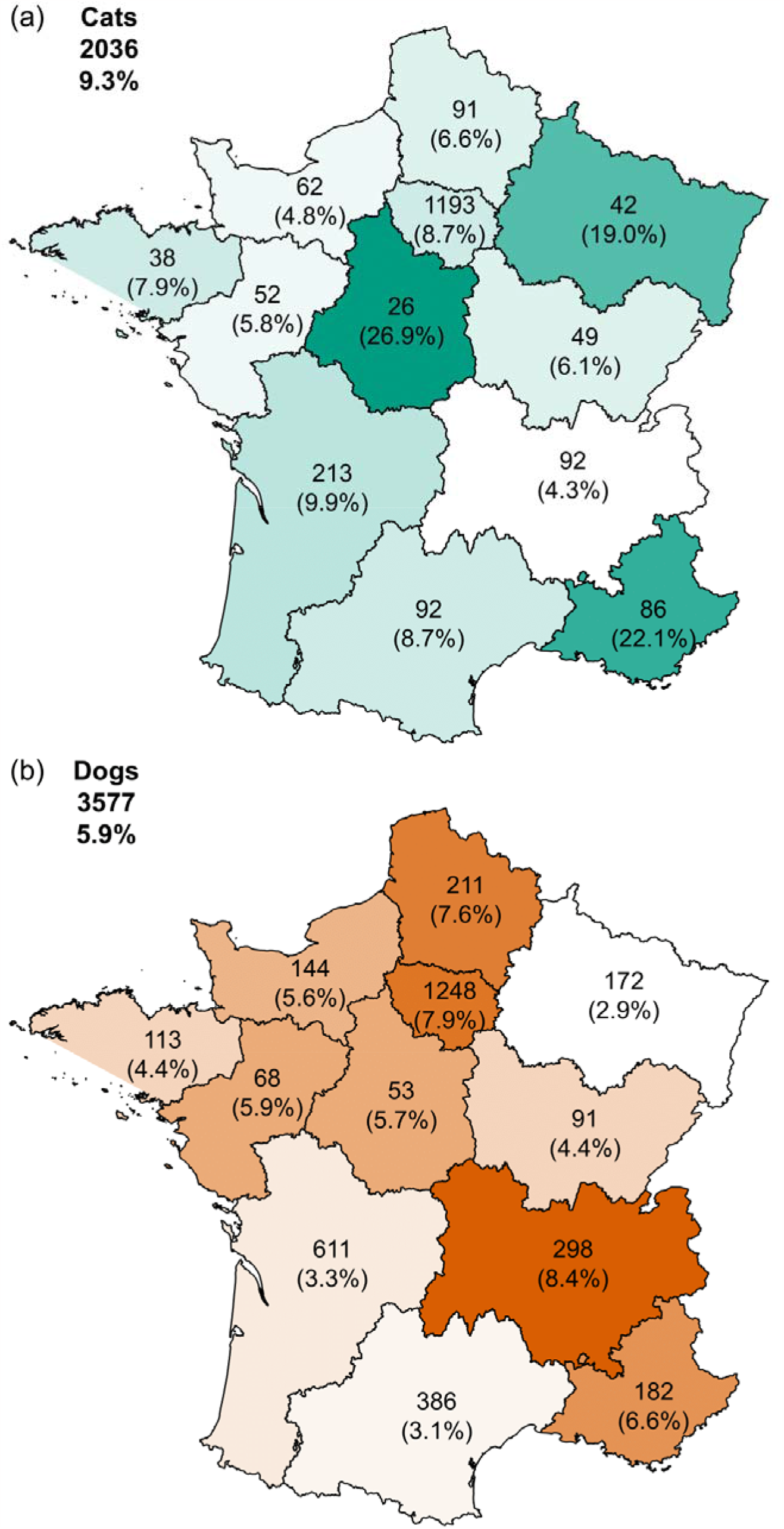
**(a). Map of France showing the number of SARS-CoV-2-positive cat sera per region**. The total number of sera samples collected per region is indicated. Seroprevalence in each region is indicated as a percentage. Regions are shaded in green according to seroprevalence. The total number of sera samples and global seroprevalence for France is in the top left corner. **(b). Map of France showing the number of SARS-CoV-2-positive dog sera per region**. The total number of sera samples collected per region is indicated. Seroprevalence in each region is indicated as a percentage. Regions are shaded in orange according to seroprevalence. The total number of sera samples and global seroprevalence for France is in the top left corner.

### Global seroprevalence

For the sera samples, 401 (7.1%) showed a positive result either against RBD, tri-S, or both (Supplementary Table 1). We next determined the presence of antibodies with neutralizing activity among the positive sera. To save time, we randomly tested approximately 75% (308) of positive sera samples. Seroneutralizing activity was detected in 81 (26.3%) of the 308 pet sera samples. Among these positive samples, 39 (48%) were positive for both RBD and tri-S, 39 (48%) were positive only for tri-s and 3 (4%) were only positive for RBD. Only the seroprevalence from MIA assays was analyzed in the remainder of the study.

### Seroprevalence in cats and dogs

We observed that a significantly greater proportion of cats were positive (189/2036, 9.3%) than dogs (212/3577, 5.9%); OR = 1.62, 95% c.i. [1.32 - 1.99], P-value = 3.8e-06, Table 2. In addition, sera from MIA-positive cats were more likely to show neutralizing activity (49/144, 34%) than dogs (32/164, 19.5%); OR = 2.12, 95% c.i. [1.27 - 3.57], P-value = 0.0039). Species differences were not always significant within each region, likely due to reduced statistical power. However, when differences were significant, it was always the case that cats were more likely to be positive than dogs. (Table 2).

**Table 2.** Seroprevalence of IgG SARS-CoV-2 antibodies detected in blood samples from cats and dogs collected in different French regions from October 2020 through June 2021. Data are presented as No. positive, percentage (95 % exact binomial confidence intervals). Odds ratios were computed by fitting binomial models region by region; an OR > 1 indicates cats were more likely to be positive than dogs. In this analysis, individual data on pet age and sex were not considered, as age was not available for all animals. P-values were computed by the likelihood ratio test.

| Region | Cats | Cats<br>seroprevalence |  | Dogs | Dogs<br>seroprevalence |  | OR (95% c.i.) | P-value |
| --- | --- | --- | --- | --- | --- | --- | --- | --- |
| Auvergne-Rhône-Alpes | 4/92 | 4.3% | (1.2-10.8) | 25/298 | 8.4% | (5.5-12.1) | 0.50 (0.17 - 1.47) | 0.17 |
| Bourgogne-Franche-Comté | 3/49 | 6.1% | (1.3-16.9) | 4/91 | 4.4% | (1.2-10.9) | 1.42 (0.30 - 6.61) | 0.66 |
| Bretagne | 3/38 | 7.9% | (1.7-21.4) | 5/113 | 4.4% | (1.5-10.0) | 1.85 (0.42 - 8.14) | 0.43 |
| Centre-Val de Loire | 7/26 | 26.9% | (11.6-47.8) | 3/53 | 5.7% | (1.2-15.7) | 6.14 (1.44 - 26.2) | 0.0098 |
| Grand Est | 8/42 | 19.0% | (8.6-34.1) | 5/172 | 2.9% | (1.0-6.7) | 7.86 (2.42 - 25.5) | 0.00057 |
| Hauts-de-France | 6/91 | 6.6% | (2.5-13.8) | 16/211 | 7.6% | (4.4-12.0) | 0.86 (0.33 - 2.27) | 0.76 |
| Île-de-France | 104/1193 | 8.7% | (7.2-10.5) | 98/1248 | 7.9% | (6.4-9.5) | 1.12 (0.84 - 1.49) | 0.44 |
| Normandie | 3/62 | 4.8% | (1.0-13.5) | 8/144 | 5.6% | (2.4-10.7) | 0.86 (0.22 - 3.37) | 0.83 |
| Nouvelle-Aquitaine | 21/213 | 9.9% | (6.2-14.7) | 20/611 | 3.3% | (2.0-5.0) | 3.23 (1.72 - 6.09) | 0.00037 |
| Occitanie | 8/92 | 8.7% | (3.8-16.4) | 12/386 | 3.1% | (1.6-5.4) | 2.97 (1.18 - 7.49) | 0.028 |
| Provence-Alpes-Côte d'Azur | 19/86 | 22.1% | (13.9-32.3) | 12/182 | 6.6% | (3.5-11.2) | 4.02 (1.85 - 8.73) | 0.00036 |
| Pays de la Loire | 3/52 | 5.8% | (1.2-15.9) | 4/68 | 5.9% | (1.6-14.4) | 0.98 (0.21 - 4.58) | 0.98 |
| **Total** | **189/2036** | **9.3%** | **(8.0 - 10.6)** | **212/3577** | **5.9%** | **(5.2 - 6.8)** | **1.62 (1.32 - 1.99)** | **3.8x10<sup>-6</sup>** |

### Seroprevalence by sex

We found no significant sex differences in seropositivity rates, either for all animals (females: 6.9%; 163/2361; males 7.5%; 212/2842; p = 0.24) or among cats (females 9.4%; 78/827, males 9.8%; 99/1009, p = 0.68) and dogs (females: 5.5%; 85/1534, males: 6.2%; 113/1833; p = 0.27) tested separately (Table 3).

**Table 3.** Seroprevalence of IgG SARS-CoV-2 antibodies in blood samples from cats and dogs by sex from October 2020 through June 2021 Data are presented as No. positive, percentage (95 % exact binomial confidence intervals). Odds ratios > 1 indicate males are more likely to be positive than females and were computed by fitting binomial generalized linear models, with age as a controlling factor. P-values correspond to likelihood ratio tests.

| Sex | Cats | Cat seroprevalence | Dogs | Dog seroprevalence | Cats + Dogs | Cats + Dogs seroprevalence |
| --- | --- | --- | --- | --- | --- | --- |
| Female | 78/827 | 9.4% (7.5-11.5) | 85/1534 | 5.5% (4.4-6.8) | 163/2361 | 6.9% (5.9-8.0) |
| Male | 99/1009 | 9.8% (8.0-11.8) | 113/1833 | 6.2% (5.1-7.4) | 212/2842 | 7.5% (6.5-8.5) |
| **Total** | **177/1836** | **9.6% (8.3-11.1)** | **198/3367** | **5.9% (5.1-6.7)** | **375/5203** | **7.2% (6.5-7.9)** |
| OR | 1.08 (0.75 - 1.54) |  | 1.20 (0.87 - 1.67) |  | 1.15 (0.91 - 1.47) |  |
| P-value | 0.68 |  | 0.27 |  | 0.24 |  |

### Seroprevalence by age

Age was reported for 1657 cats (range: 0.2 – 22yr) and 2781 dogs (range: 0.1 – 18.5yr). Among cats, 18.4% aged [0-3] years, 10.4% aged]3-9] years, and 6.4% aged over 9 years tested positive. Among dogs, 8.8% aged [0-3] years, 5.8% aged]3-9] years, and 5.1% aged over 9 years tested positive. Using a binomial model with age entered as a continuous variable, we observed a significant decrease in seroprevalence with age in cats (OR for a one-year increase in age = 0.91, 95% c.i. [0.88 - 0.94], p-value = 3.7e-08) and dogs (OR = 0.95, 95% c.i. [0.92 - 0.99], p-value = 0.016) (Figure 3) (Supplementary Table 2).

**Figure 3.**
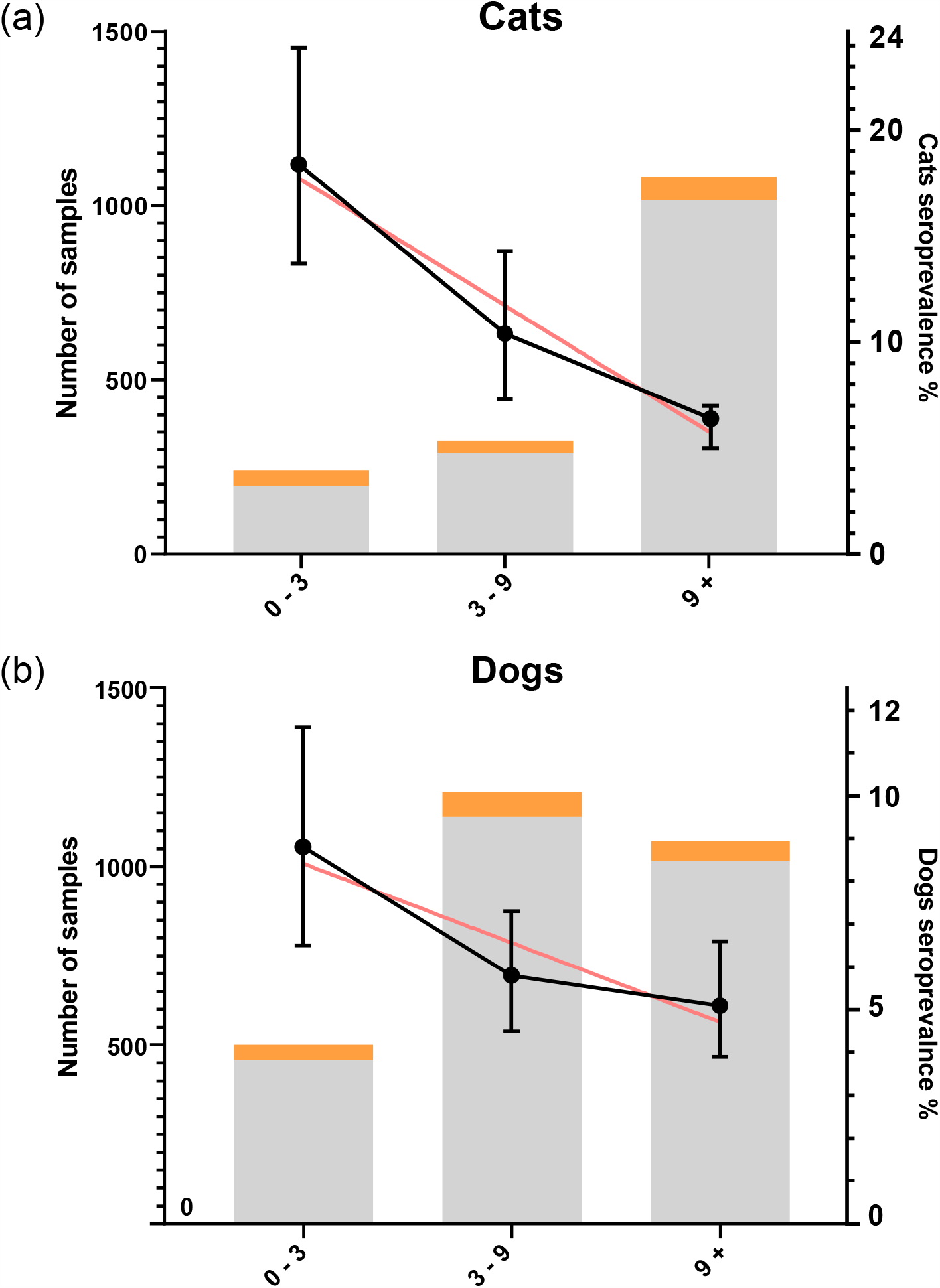
**(a). The number of cat blood samples tested by age group for anti-SARS-CoV-2 antibodies by MIA from October 2020 through June 2021**. Samples testing negative are shaded grey, and seropositive samples are in orange. Seroprevalence is represented by black dots, with 95 % binomial confidence interval. The red line represents the linear regression. **(b). The number of dog blood samples tested by age group for anti-SARS-CoV-2 antibodies by MIA from October 2020 through June 2021**. Samples testing negative are shaded grey, and seropositive samples are in orange. Seroprevalence is represented by black dots, with 95 % binomial confidence intervals. The red line represents the linear regression.

### Seroprevalence over time

We next examined whether seroprevalence was associated with the time of sampling. For this analysis, we selected animals at least one year old at the date of sampling (Figure 4). Seroprevalence was not associated with the time of sampling for cats: p-value = 0.41. However, seroprevalence among dogs increased over the 9 months of the study (OR = 3.47, 95% c.i. [1.47 - 8.23], p-value = 0.0045).

**Figure 4.**
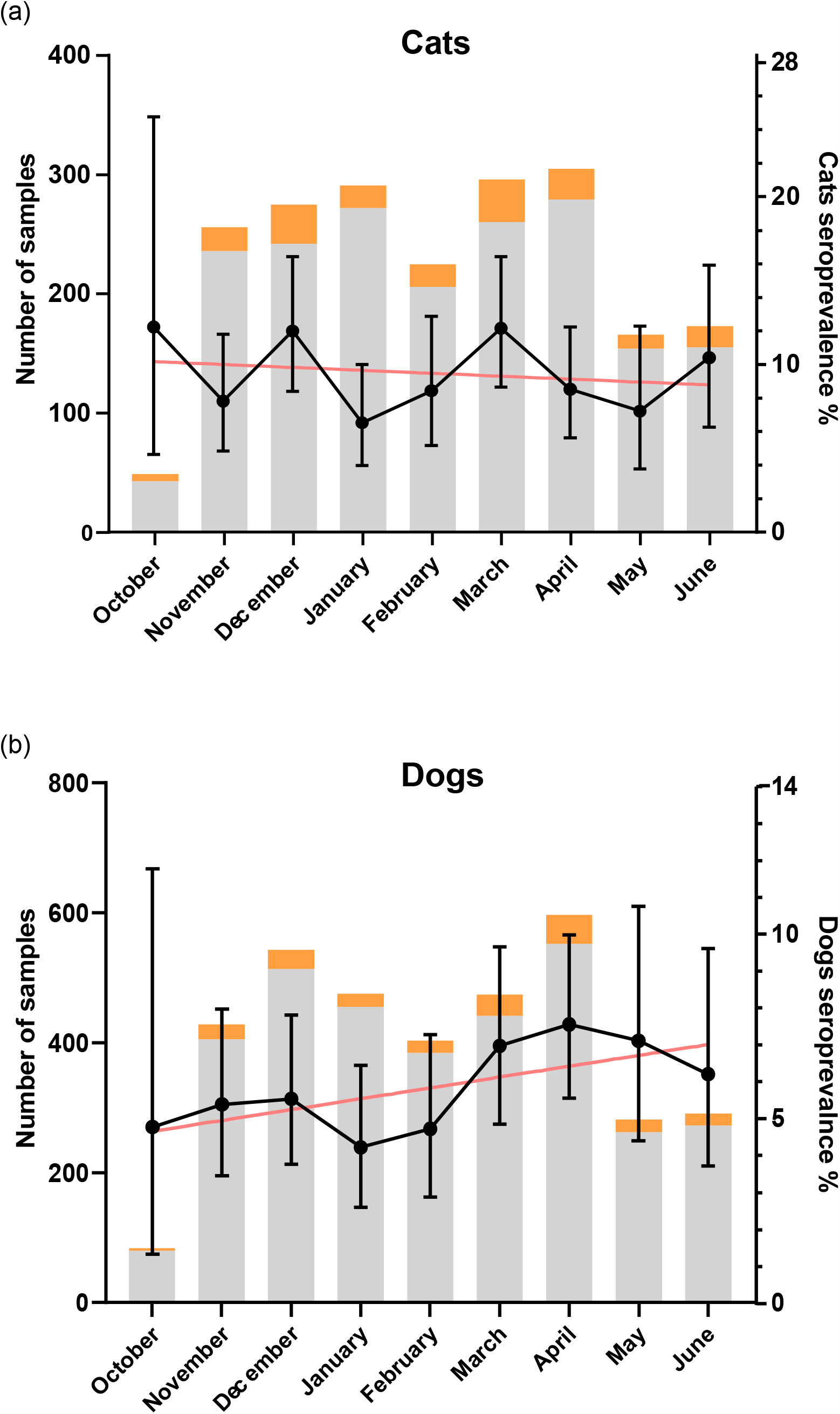
**(a). The number of cat blood samples tested each month for anti-SARS-CoV-2 antibodies by MIA from October 2020 through June 2021**. Samples testing negative are shaded grey, and seropositive samples are in orange. Seroprevalence is represented by black dots, with 95 % binomial confidence interval. The red line represents the linear regression. **(b). The number of dog blood samples tested each month for anti-SARS-CoV-2 antibodies by MIA from October 2020 through June 2021**. Samples testing negative are shaded grey, and seropositive samples are in orange. Seroprevalence is represented by black dots, with 95 % binomial confidence interval. The red line represents the linear regression. Notice that in this figure, dates have been pooled by calendar month for illustrative purposes but that in the statistical analysis exact dates were used. Likewise, the regression lines are similarly illustrative as the statistical tests were based on logistic (binomial) regression.

## Discussion

This study reports a large-scale serological survey of pet (cats and dogs) to detect anti-SARS-CoV-2 IgG antibodies. The samples were collected in metropolitan France from October 2020 through June 2021, a period including peaks of the second and third waves of SARS-CoV-2 infections in France. From a sample of 5613 pets, we reported a seroprevalence of anti-SARS-CoV-2 antibodies of 7.1%. We observed that only a small percentage of samples (48%) were positive for both tri-s and RBD, indicating that the RBD assay may be less sensitive than the tri-s assay. This may be explained by the fact that the full trimeric spike antigen may bind a broader range of antibodies than the receptor-binding domain, which includes only a small part of the spike protein. We found neutralizing antibody activity in the sera of only 26% of seropositive pets. Previous studies have shown that some pets do not develop neutralizing antibodies (5, 30). Cats were more likely to produce neutralizing antibodies than dogs, which is likely associated with a more prolonged and intense immune stimulation in cats. In humans, disease severity is positively correlated with neutralizing antibody levels(31).

In cats, we found a higher seroprevalence (9.3%) than previously observed in other European countries, which ranged from 0% to 6.4% (13, 17-19, 21-24). However, most of these studies were done before the second wave, during a period of relatively lower viral circulation than our sampling period. In addition, most of these studies used a seroneutralisation assay.

In dogs, the observed seroprevalence (5.9%) is in accord with a previous study in France showing a prevalence of 4.8% in companion and military working dogs sampled between February 2020 and February 2021 (16). Other studies looking for SARS-CoV-2 antibodies in dogs have reported seroprevalences ranging from 0% to 14.5% (10, 13, 17-19, 21).

Importantly, we observed a significantly higher seroprevalence of anti-SARS-CoV-2 antibodies in cats than in dogs (*p* = 4.2e-08). The statistical significance of this difference varied among regions, likely due to the reduced power and perhaps some unintended sampling bias by veterinarians. For example, the smallest sample size was in the Bourgogne-Franche-Comté region, where we observed no significant difference between dogs and cats. Furthermore, for a region like Ile-de-France, where people live mostly in apartments, we can also hypothesize that dogs live in closer contact with owners than in the rest of France. Previous studies with fewer samples have either found no significant difference between species (8, 9, 13, 17, 19, 32) or that cats have significantly higher seroprevalence than dogs (3, 6, 7).

Our study of a very large population of dogs and cats in natural conditions provides some evidence that cats are more susceptible to SARS-CoV-2 infection than dogs, at least during the time frame of our sampling period. Potential causes of species differences in susceptibility between cats and dogs are numerous, but likely include a variety of biological and behavioural factors, as well as differences in exposure. Intererestingly, ACE-2 shows greater sequence similarity between cat and human orthologs than observed between dogs and humans (33). The absence of data such as the pet lifestyle (Indoor/Outdoor), or the frequency and nature of contacts with humans and other animals restricts our ability to identify a potential cause of the observed difference. In previous studies, most infected pets were epidemiologically linked to humans who had tested positive for COVID-19 (34).

We did not observe significant sex differences in seroprevalence in either species (*p* = 0.45). Our findings are consistent with most previous studies also reporting an absence of sex differences in dogs and cats (6, 17, 32, 35). A smaller study of 188 dogs and 61 cats found higher seropositivity in male dogs and an absence of a sex difference in cats (9). Another study found that male dogs sampled from the general population were more likely to test positive than females, but this difference was not observed in dogs from COVID-19+ households (10). There is little evidence of a significant sex difference in susceptibility in humans. However, men are more likely to be affected by severe forms of COVID than women for a variety of reasons (36).

In terms of age, we observed a higher seroprevalence among younger animals (between 0-3 years) for both species that then decreased with age. A study of dogs sampled from the general population found seroprevalence was highest in animals aged 5-6 years and that in COVID-19+ households, seroprevalence peaked in slightly younger dogs, aged between one and five years (10). Other studies have reported no significant associations with age in cats and dogs (6, 17). An experimental study in cats found that juveniles appear more vulnerable than subadults (27). The decreasing seroprevalence we observed with age could also arise from age-dependent behavioural changes. For example, young animals (< 3 years old) are more active and curious and may be in greater contact with their owners than older animals that prefer to remain quieter. The decrease could also reflect immunosenescence in older animals, as observed in humans.

Interestingly, we observed a slight increase in seroprevalence in dogs during the study’s nine months of sampling, a trend not observed among cats. We expected an increase because antibodies have a longer persistence in the organism than viral RNA; thus, animals sampled at later dates would represent an accumulation of cases. The absence of a positive association between seroprevalence and the time of sampling in cats has been reported in two other studies in Europe, but conclusions were limited by the small number of samples collected over just a few months (24, 37). The absence of an association in cats suggests a limited persistence of antibodies in cats than dogs. Few studies have investigated variation in the persistence of antibodies in animals. For example, a study carried out on seven dogs and two cats infected in natural conditions showed persistence of neutralizing antibodies up to 10 months after infection in four of the dogs and the two cats, but also that persistence was markedly reduced in two of the dogs after three months (38). Moreover, a study of two cats found that neutralizing antibodies had disappeared by 110 days (25). Based on these data, one possible reason for the lack of increase in seroprevalence during our study period could be a progressive seroreversion of infected cats that is equally compensated by the number of new infections, i.e. seroconversion. If so, this would mean that the observed seroprevalence is not an accurate reflection of the total number of infections, at least in cats, during the whole epidemic. Instead, seroprevalence provides a snapshot if infections acquired during a time period that remains to be defined by longitudinal serological studies of cats and dogs. This also suggests that the seroprevalence observed in our study may underestimate the actual proportion of cats infected during the entirety of the epidemic.

Human-to-pet transmission may promote viral adaptation facilitating re-infection with novel viral strains in humans (39). While one case of infection from cat to human has recently been reported, the large number of pet cats and their frequent close interaction with humans provides ample opportunity. This possibility raises the question of a vaccination strategy for animals susceptible to SARS-CoV-2 infection. While pets do not currently seem to play a role in the ongoing pandemic, our results emphasize the magnitude of SARS-CoV-2 infection in pets is not trivial. Combined with the size of domestic cat and dog populations and the close contact with their human companions, our results highlight the importance of collecting more data on SARS-CoV-2 transmissibility and pathogenicity in companion animals, especially with the emergence of new variants. Also, when a SARS-CoV-2 infection is suspected in a pet, we suggest collecting a sample for RT-qPCR confirmation of infection, followed by whole-genome sequencing to identify new mutations, particularly in antigenic sites targeted by the immune system. Finally, similar public health recommendations applied to humans should also be implemented for animals to prevent human-to-animal transmission, such as not having contact with animals when a household member is COVID-19 positive.

## Acknowledgements

We are grateful to the pet owners for giving us permission to sample their pets. We thank the veterinarians that helped us with sampling. We thank Simon Thierry and Trung Thanh Nguyen for technical support and assistance. We also thank Kurt McKean (https://octopusediting.com/) for the English editing of the manuscript.

## Institutional Review Board Statement

According to the act of “use of live animals for scientific purposes” effective in France on 14 January 2022, ethical approval was not sought or required since all pets were sampled by a veterinarian during a health care visit. All applicable international and national guidelines for the care of pets were followed.

## Informed Consent Statement

Not applicable.

## Data Availability Statement

Raw data and scripts are available on zenodo: https://doi.org/10.5281/zenodo.7646559

## Author Contributions

M.F., P.B., S.G.R., A.B.-M., V.L., and E.M.L. conceived and designed the study.

M.F., E.E., D.d.R.d.F., D.G., S.D., B.B., and V.L designed and performed the experiments.

All authors analyzed the data and interpreted and discussed the results.

M.F. and E.L. wrote the manuscript with input from all authors.

All authors have read and agreed to the published version of the manuscript.

