## Supplementary Table 1 for "A large-scale serological survey in pets from October 2020 through June 2021 in France shows significantly higher exposure to SARS-CoV-2 in cats" (from another version of this preprint: version 3, 2023-07-10)

**Supplementary table 1.** Distribution of positive results in blood samples from cats and dogs by antigen from October 2020 through June 2021.

|  | RBD | S tri | RBD and S tri | **Total** |
| --- | --- | --- | --- | --- |
| Cats | 23 | 111 | 55 | **189** |
| Dogs | 20 | 134 | 58 | **212** |
| **Total** | **43** | **245** | **113** | **401** |
