## Supplementary Table 2 for "A large-scale serological survey in pets from October 2020 through June 2021 in France shows significantly higher exposure to SARS-CoV-2 in cats" (from another version of this preprint: version 3, 2023-07-10)

**Supplementary table 2.** Seroprevalence of IgG SARS-CoV-2 antibodies in blood samples from cats and dogs by age from October 2020 through June 2021. Based on seroprevalence from MIA.

| Age class | Cats | Cat prevalence | | Dogs | Dog prevalence (%) | |
| --- | --- | --- | --- | --- | --- | --- |
| 0 - 3 | 44/239 | 18.4% | (13.7 – 23.9) | 44/501 | 8.8% | (6.5 -11.6) |
| 3 - 9 | 34/326 | 10.4% | (7.3 – 14.3) | 70/1209 | 5.8% | (4.5 – 7.3) |
| 9 + | 69/1084 | 6.4% | (5.0 – 7.0) | 55/1071 | 5.1% | (3.9 – 6.6) |
| **Total** | **147/1657** | **8.9%** | **(7.5 – 10.3)** | **169/2781** | **6.1%** | **(5.2 – 7.0)** |

Data are presented as No. positive, percentage (95 % exact binomial confidence intervals). P-values correspond to the Fisher exact test comparing seroprevalence in cats vs dogs
